# A database of egg size and shape from more than 6,700 insect species

**DOI:** 10.1101/471953

**Authors:** Samuel H. Church, Seth Donoughe, Bruno A. S. de Medeiros, Cassandra G. Extavour

**Author notes:** Samuel H. Church and Seth Donoughe contributed equally to this work.

## Abstract

Offspring size is a fundamental trait in disparate biological fields of study. This trait can be measured as the size of plant seeds, animal eggs, or live young, and it influences ecological interactions, organism fitness, maternal investment, and embryonic development. Although multiple evolutionary processes have been predicted to drive the evolution of offspring size, the phylogenetic distribution of this trait remains poorly understood, due to the difficulty of reliably collecting and comparing offspring size data from many species. Here we present a database of 10,449 morphological descriptions of insect eggs, with records for 6,706 unique insect species and representatives from every extant hexapod order. The dataset includes eggs whose volumes span more than eight orders of magnitude. We created this database by partially automating the extraction of egg traits from the primary literature. In the process, we overcame challenges associated with large-scale phenotyping by designing and employing custom bioinformatic solutions to common problems. We matched the taxa in this database to the currently accepted scientific names in taxonomic and genetic databases, which will facilitate the use of this data for testing pressing evolutionary hypotheses in offspring size evolution.

## 2 Background & summary

The size of a reproductive propagule, for example an animal egg or a plant seed, has crucial implications for the biology of both the parent and the offspring^1–3^. From the perspective of the parent organism, propagule size is a component of the maternal investment in each offspring^2^, and propagule size is predicted to be positively correlated with adult body size and negatively correlated with propagule number^3–5^. From the perspective of the offspring, the size of the propagule is relevant to the starting material for embryonic development, and it can impact both life history and ecological interactions^2,6^. Evolutionary hypotheses have been proposed to explain patterns in the diversity of propagule size, yet the robustness or generality of the patterns themselves have rarely been tested across species^3^. To understand the evolutionary forces driving propagule size evolution, we need large-scale, reliable descriptions of the distribution of propagule size across the evolutionary tree.

Insect eggs come in an incredible diversity of shapes and sizes^7,8^. The thousands of egg descriptions in the entomological literature, however, have never to our knowledge been systematically compiled across insects. Without a comparison of egg sizes across insects, we cannot ascertain basic information such as the extant range of insect egg sizes, or the relationship between size and ecology or development. To address this problem, we created a database of quantitative parameters describing egg morphology from the entomological literature. All data were collected from published records, including both measurements reported in text descriptions of insect eggs, as well as our own new measurements of published images. We developed custom software that allowed us to collect data from thousands of publications efficiently and reproducibly (Figure 1). We provide this software as a set of tools that can assist other scientists in collecting phenotypic data from the literature (see Methods).

Using this software we extracted egg descriptions from 1,756 publications from the past 250 years (Table 1). The database has 10,449 entries representing every extant order of insects, and 6,706 unique insect species. The insect egg database includes descriptions of egg size and shape (Table 2), and the scientific name of each entry has been matched to current taxonomic and genetic databases. The egg database is made publicly available for download (see Methods).

**Table 1:** Bibliographic, data type, and taxonomic statistics of the insect egg database

| **Bibliographic statistics** |  |
| --- | --- |
| references examined | 2900 |
| references with egg information | 1756 |
| unique authors | 1498 |
| unique journals / books | 491 |
| **Data type statistics** |  |
| Total entries in egg database | 10449 |
| Entries with text description of length and width | 7672 |
| Length reported as average and deviation | 1065 |
| Length reported as range | 2188 |
| Single length value reported | 4419 |
| Only volume reported | 1368 |
| Entries with an image | 4774 |
| Images re-measured | 2004 |
| Entries with both text and image measurements | 1205 |
| **Taxonomic statistics** |  |
| unique hexapod species | 6706 |
| unique hexapod genera | 4077 |
| unique hexapod families | 526 |
| unique hexapod orders | 32 |

**Table 2:** Trait def initions and standardizations. * volume was included only when length and width measurements were not available from text. ** measurements included only when a scale bar was published with the image.

| Text measurements |  | Standardized text measurements |  |  |
| --- | --- | --- | --- | --- |
| Name | Units | Name | Units | Method |
| length or height | mm | length, $l$ | mm | as recorded |
| width or diameter | mm | width, $w$ | mm | $\max(w, b)$ |
| breadth or depth | mm | breadth, $b$ | mm | $\min(w, b)$ |
| volume* | mm<sup>3</sup> | volume, $v$ | mm<sup>3</sup> | $\frac{1}{6}\pi lwb$ OR $\frac{1}{6}\pi lw^2$ OR $v$ |
| | | aspect ratio | ratio, no units | $\frac{l}{w}$ |

| Image measurements |  | Standardized image measurements |  |  |
| --- | --- | --- | --- | --- |
| Name | Units | Name | Units | Method |
| curved length | px | length**, $l$ | mm | straight length |
| straight length | px | width**, $w$ | mm | $\max(q_1, q_2, q_3)$ |
| 1st quartile width, $q_1$ | px | volume** | mm<sup>3</sup> | $\frac{1}{6}\pi lw^2$ |
| 2nd quartile width, $q_2$ | px | aspect ratio | ratio, no units | $\frac{l}{w}$ |
| 3rd quartile width, $q_3$ | px | asymmetry | ratio, no units | $\frac{\max(q_1, q_3)}{\min(q_1, q_3)} - 1$ |
| angle of curvature | degrees, radians | angle of curvature | radians | as recorded |

| Final database measurements |  |  |  |
| --- | --- | --- | --- |
| Name | Units | Transformation | Method |
| length | mm | $\log_{10}$ | used text measurement, when both text and image were available |
| width | mm | $\log_{10}$ | used text measurement, when both text and image were available |
| breadth | mm | $\log_{10}$ | used text measurement, when both text and image were available |
| volume | mm<sup>3</sup> | $\log_{10}$ | used text measurement, when both text and image were available |
| aspect ratio | ratio, no units | $\log_{10}$ | used text measurement, when both text and image were available, removed egg images in the top 0.1% |
| asymmetry | ratio, no units | sq. root | removed egg images in the top 0.1% |
| angle of curvature | radians | sq. root | did not record for eggs with an aspect ratio $\leq 1$ |

Insect egg sizes vary between species, within species, and within a single individual^7^, and the database described here contains variation from all of these sources. We calculated the degree of intraspecific variation in egg length for all taxa where these data were available in the literature. We additionally assessed the variation in the precision used to record data for all database entries. This provides the necessary information to account for sources of variation in a comparative study of insect egg morphology.

The insect egg database includes representatives of all insect orders (Table 1), but these orders are not equivalent to each other either in terms of number of extant species or in the historical degree of entomological study^9,10^. We therefore assessed the phylogenetic coverage of the insect egg database relative to the number of species estimated for each clade. This enables evaluation of the potential bias present in the database, and highlights undersampled clades as potential priorities for future study.

The methods used to create the insect egg database include solutions to challenges in assembling phenotypic data from large groups of organisms. Phenotypic descriptions can require great resources and expertise to reliably collect, identify, and describe morphological features across thousands of species^11^. This expense can limit macroevolutionary studies of morphological evolution. One way to overcome this barrier is to rely on the thousands of data points already reported by experts in the scientific literature. However, this method brings its own challenges, such as assigning concordance between taxonomic names and extracting data from published text or images^11^. To address these needs, we include bioinformatic approaches that can be used by future researchers. Both the egg database and the software solutions used to generate it will have broad value for researchers interested in studying questions of morphological evolution across large evolutionary scales.

## 3 Methods

### 3.1 Gathering primary literature with egg descriptions

The workflow used to assembling the database is shown in Figure 1. Publications were identified for potential inclusion in the egg database using the following online literature databases: Google Scholar (scholar.google.com), Web of Knowledge (webofknowledge.com), and Harvard‘s HOLLIS library system (hollis.harvard.edu). We searched these databases continuously during the period of from October 2015 – August 2017 with a predetermined set of word pairs that included an insect common or taxonomic name (e.g. ‘fly’, ‘Diptera’, ‘Nematocera’) and one of the following egg related terms: ‘egg’, ‘chorion’, ‘immature’, or ‘embryo’. Insect clade names included all insect order names and all insect families from the five largest insect orders (Coleoptera, Diptera, Lepidoptera, Hymenoptera, and Hemiptera). Following a search, all publications returned by the search were manually evaluated for inclusion in the database. The criteria for this evaluation were as follows: [1] Does the title or abstract of the paper suggest that the paper contains insect egg information? [2] If the publication could be immediately previewed on the Harvard library system, does it contain an egg measurement in the text or an egg image with a scale bar? [3] If the publication could not be immediately previewed, does the title or abstract refer to descriptions of the chorion, immature stages, or embryology? If a publication met at least one of these criteria, complete bibliographic information for the reference was stored in a master BibTeX reference file (available at Dryad https://datadryad.org/review?doi=doi:10.5061/dryad.pv40d2r). Publications were continually added to the database throughout the study, and the final count of publications that met these criteria were 2,900, of which 1,756 contained egg morphological data. The language of the publication was not a criterion for inclusion in the database. However, due to the nature of the online search engines that we used, the database is enriched for papers published with at least an abstract in English. A formatted list of the references cited in the egg database is available in the supplemental file ‘bibliography_egg_database’.

**Figure 1:**
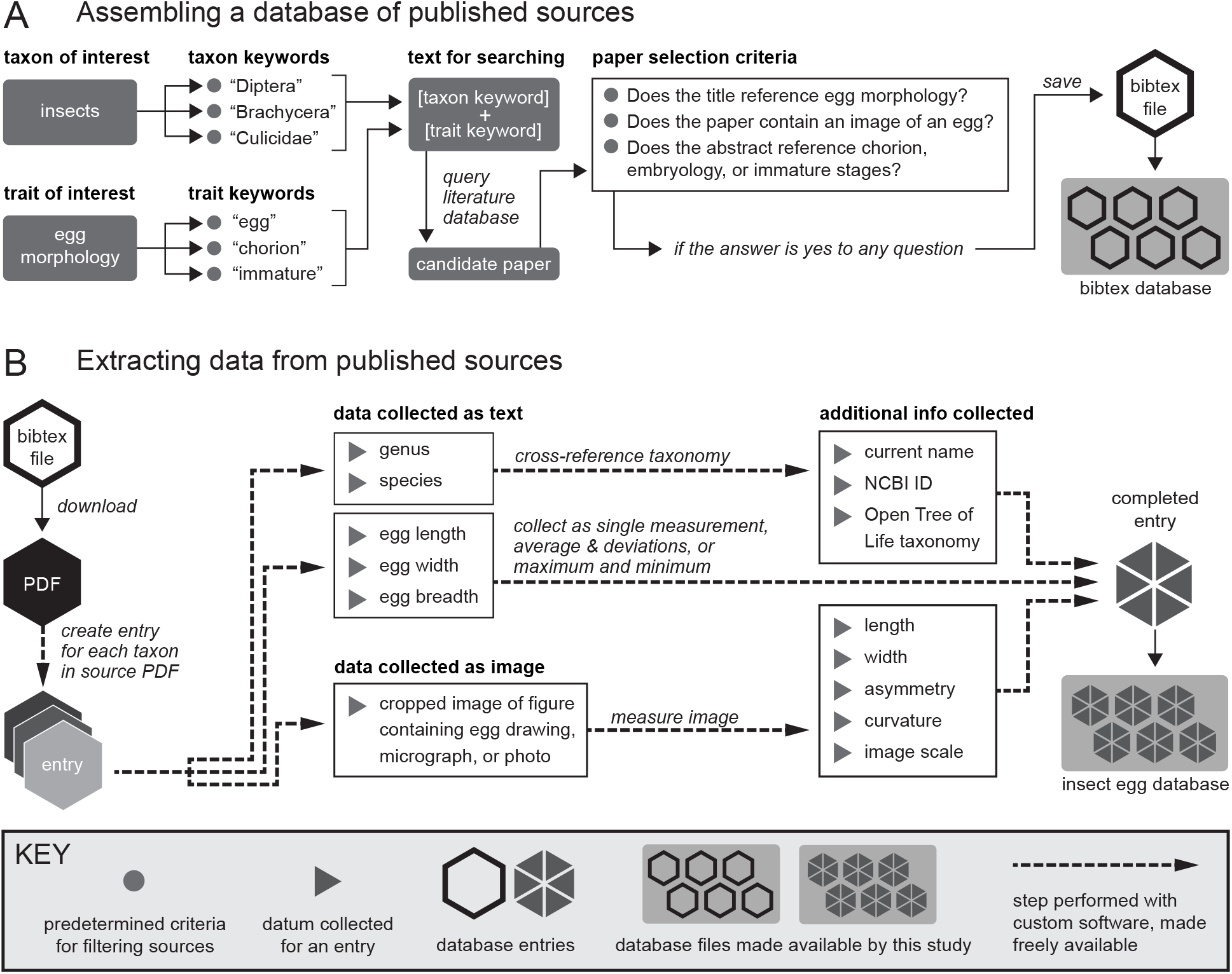
The workflow used to create the insect egg database. The database was compiled from the insect literature following the discrete steps shown here, using custom bioinformatic software to maximize reproducibility, consistency, and efficiency. **A**, The workflow used to evaluate candidate publications for inclusion in the database. **B**, The workflow used to extract egg descriptions from the text of published sources and to remeasure published images of eggs. Steps performed with custom software are shown in dashed lines.

### 3.2 Def ining egg traits

The egg traits in the database are listed in Table 2. For each trait listed below we used the descriptions of egg length and width as presented in the original publications. Given that conventions vary across entomologists and insect taxonomic groups, we present the following definitions to resolve ambiguous cases and to serve as a suggestion for future egg descriptions.

#### Egg

The term *egg* is used in the literature to describe several successive developmental stages, including the mature oocyte, the zygote cell, and the developing embryo in its eggshell. For consistency we selected measurements that were recorded closest to the time of fertilization, when multiple descriptions were available within a single publication, given that in some insects it has been documented that the dimensions of the egg change over time (typically <20% change in length due to water exchange during embryonic development)^7,12–15^. In most insects the egg is oviposited outside the adult body; however in viviparous insects, eggs proceed through some or all of embryonic development within the body of the mother. The egg is often enveloped in a secreted eggshell called the chorion^15^, which may have elaborations (e.g. dorsal appendages or opercula)^16^. We selected egg measurements that excluded chorionic elaborations over those that included them, as our goal was to measure the comparable cellular material across species.

#### Length

To resolve ambiguous cases, and when measuring egg features from published images, we defined egg length as the distance in millimeters (mm) of the axis of rotational symmetry. This definition maximizes consistency with published descriptions of egg length. Under this definition, length is not always longer than width (as defined below). For some insect groups (e.g. Lepidoptera) the axis of rotational symmetry is sometimes referred to in the literature as *height*^17–19^. For published images with a scale bar, we measured both the straight and curved length of the egg (for those eggs that are curved), but for all analyses and figures, we used the straight length of the egg to maximize consistency with published records.

#### Width and breadth

To resolve ambiguous cases, and when measuring egg features from images, we defined width as the widest diameter (mm) measured perpendicular to the axis of rotationally symmetric axis of the egg. For some insect groups this axis is referred to in the literature as *diameter*^17^ or *breadth*^20^. For eggs described in published records as having a length, width, and breadth or depth (i.e., the egg is a flattened ellipsoid^21^), we considered *width* as the wider of the two diameters, and *breadth* as the diameter perpendicular to both width and length. For published images with a scale bar, we measured width as the widest of the three egg diameters at the first quartile, midpoint, and third quartile of the length axis. We did not measure breadth from published images.

#### Volume

Volume (mm^3^) was calculated using the equation for the volume of an ellipsoid, following previous studies^22,23^. The formula is 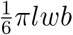, with *l, w*, and *b* as length, width, and breadth, respectively. This simplifies to 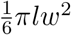 when the egg is rotationally symmetric. For records in which the volume was reported but egg length and width were not, we used the reported volume. For all other entries, we recalculated volume from the measurements in the text and from measurements of images published with a scale bar.

#### Aspect ratio

We calculated aspect ratio as the ratio of length to width. An aspect ratio of one corresponds to a spherical egg. An aspect ratio less than one corresponds to an egg that is wider than long (oblate ellipsoid). An aspect ratio greater than one corresponds to an egg that is longer than it is wide (prolate ellipsoid). Analyses testing the sensitivity of our measurement software (see “Assessing the accuracy of image measuring software” below) for egg images indicated that the variance in measured aspect ratio increases sharply when aspect ratio is much higher than typical (Table 3). Therefore we excluded the eggs in the top 0.1 percentile of aspect ratio from the final database. We recorded the aspect ratio from images published with or without a scale bar, as aspect ratio is a scale-free attribute.

**Table 3:** Results of image measurement software accuracy assessment. Mean discrepancy calculated as the average difference between the actual and measured values, n = 5.

| Aspect ratio | Actual value |  | Aspect ratio | Mean discrepancy |  |
| --- | --- | --- | --- | --- | --- |
|  | Asymmetry | Angle of curvature |  | Asymmetry | Angle of curvature |
| 0.5 | 0 | 0 | -0.01 | -0.05 |  |
| 0.5 | 0.2 | 0 | -0.01 | -0.08 |  |
| 0.5 | 0.8 | 0 | -0.02 | 0.02 |  |
| 1 | 0 | 0 | -0.02 | -0.05 |  |
| 1 | 0.2 | 0 | -0.03 | -0.07 |  |
| 1 | 0.8 | 0 | -0.03 | -0.13 |  |
| 2 | 0 | 0 | -0.03 | -0.04 | -2.68 |
| 2 | 0 | 30 | -0.06 | -0.04 | 8.74 |
| 2 | 0 | 120 | -0.18 | -0.05 | 15.49 |
| 2 | 0.2 | 0 | -0.06 | -0.05 | -2.99 |
| 2 | 0.2 | 30 | -0.05 | -0.07 | 6.66 |
| 2 | 0.2 | 120 | -0.17 | -0.02 | 16.75 |
| 2 | 0.8 | 0 | -0.09 | -0.08 | -0.65 |
| 2 | 0.8 | 30 | -0.10 | -0.14 | 15.02 |
| 2 | 0.8 | 120 | -0.18 | -0.06 | 23.84 |
| 6 | 0 | 0 | -0.36 | -0.06 | -1.63 |
| 6 | 0 | 30 | -0.15 | -0.04 | -1.47 |
| 6 | 0 | 120 | -0.32 | -0.05 | 2.52 |
| 6 | 0.2 | 0 | -0.24 | -0.06 | -0.66 |
| 6 | 0.2 | 30 | -0.50 | -0.19 | -0.80 |
| 6 | 0.2 | 120 | -0.45 | -0.06 | 3.32 |
| 6 | 0.8 | 0 | -0.36 | -0.25 | -2.61 |
| 6 | 0.8 | 30 | -0.56 | -0.13 | -0.16 |
| 6 | 0.8 | 120 | -0.40 | -0.14 | 2.28 |

#### Asymmetry

We defined asymmetry as 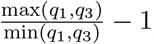, where *q*1 and *q*3 are the egg diameters at the first and third quartile of the curved length axis. Therefore an egg with an asymmetry of zero has quartile diameters with equal length. Baker’s λ value, used to measure asymmetry in bird eggs^24^, can be converted to the asymmetry parameter used in the present study. Analyses testing the sensitivity of our image measuring software (see “Assessing the accuracy of image measuring software” below) indicated that the variance increases sharply near the extreme high values of asymmetry (Table 3). We therefore excluded the eggs in the top 0.1 percentile of asymmetry from the final database. Asymmetry was only recorded from published egg images.

#### Angle of curvature

We defined the angle of egg curvature as the angle of the arc (measured in degrees) created by the endpoints and midpoint of the length axis. Analyses testing the sensitivity of our image measuring software (see “Assessing the accuracy of image measuring software” below) indicated that the variance in curvature increases when the curvature and aspect ratio are low (Table 3). We therefore did not calculate curvature for eggs with an aspect ratio of one or less. Angle of curvature was only recorded from published egg images.

### 3.3 Extracting egg descriptions from text sources

Information was extracted from publications using a custom text parsing tool that automatically opened and searched the text of a PDF of the publication (https://github.com/shchurch/Insect_Egg_Evolution, file ‘parsing_eggs.py’, commit bd765c8). The tool, written in Python2, uses a text scoring formula to identify candidate blocks of text that contain egg descriptions and corresponding names. Each database entry was manually verified and stored in tab delimited format.

All entries included, at a minimum, a genus name and an egg measurement in one dimension or egg volume. Measurements were recorded as either an average and deviation, a range of measurements, or a single value, with precedence for inclusion given in that order. A text description of the volume of the egg was included only in cases in which there were no available data on the linear dimensions of the egg. The majority of the descriptions are reported as single values (Table 1).

### 3.4 Measuring published images of eggs

Published images of eggs were measured using a custom tool (https://github.com/sdonoughe/Insect_Egg_Image_Parser, commit faee2e8) that enabled the user to calculate aspect ratio, curvature, and asymmetry of the egg by dropping guided landmarks on the published egg image (Figure 2). If the published image included a scale bar, the program also measured the absolute length and width of the egg. The final output of this tool was combined with the corresponding text description of the egg of that species. Images were included regardless of type (e.g. light micrograph, scanning electron micrograph, drawing). However, images of low quality were excluded by manually evaluating cases where landmarks could not be placed unambiguously.

**Figure 2:**
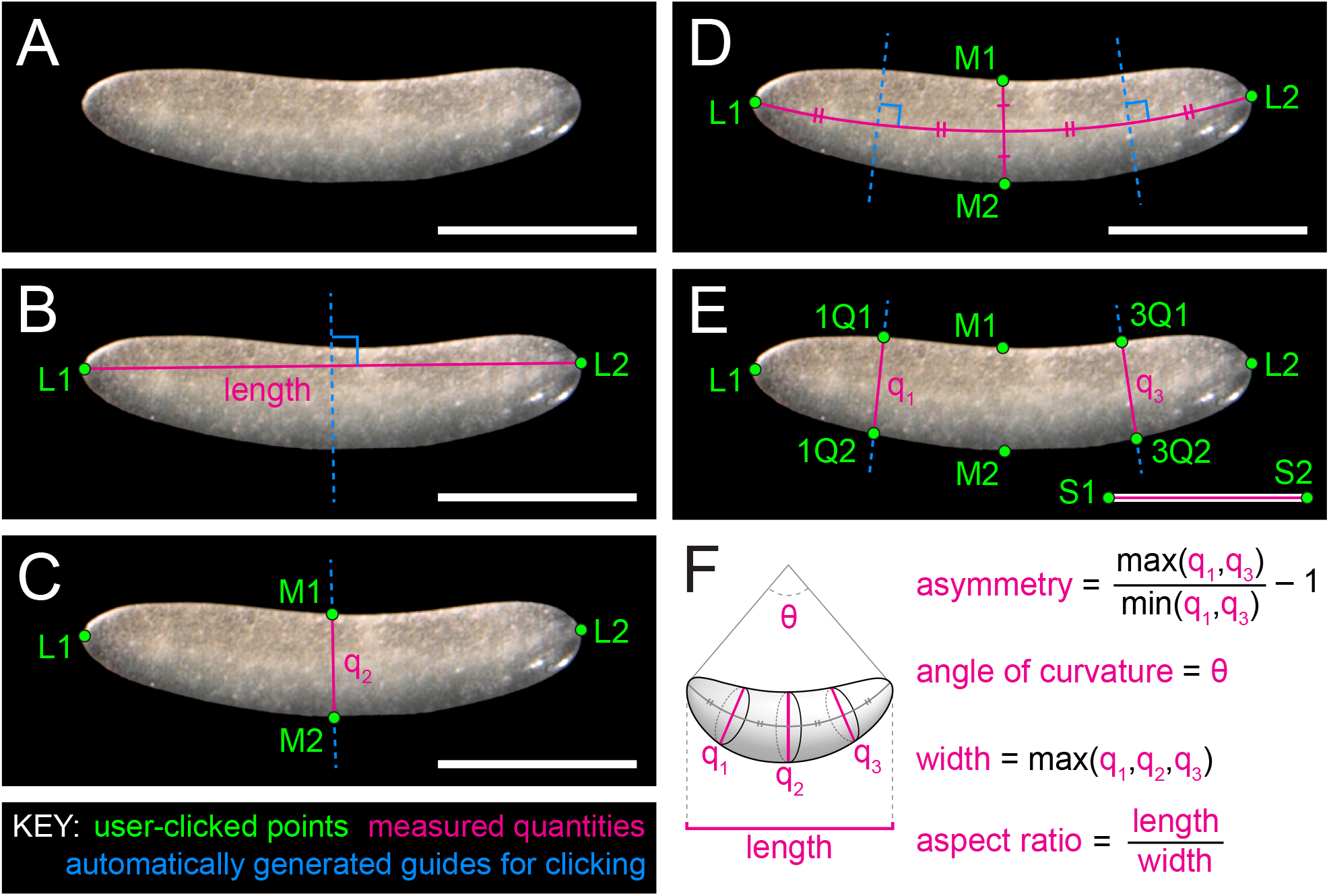
Demonstration of guided landmark-based measurement of egg shape traits. **A**, An example micrograph of an egg, in this case from the cricket *Gryllus bimaculatus*. **B**, The user places points L1 and L2 at the poles of the egg. We define egg ‘poles’ as the points on opposite sides of the egg where the curvature of the egg margin is steepest. The tool draws a line segment connecting L1 and L2 (length) and then draws its perpendicular bisector (dashed blue line). **C**, The user uses the blue line as a guide to place points M1 and M2 where the line meets the egg margin. The tool draws a line segment connecting M1 and M2 (*q*2). **D**, The tool draws a curved segment connecting the midpoint of *q*1 with L1 and L2, and then draws two perpendicular bisectors of the curved segment (dashed blue lines). **E**, The user uses the blue lines as a guide to place points 1Q1, 1Q2, 3Q1, and 3Q2 where the lines meet the egg margin. The tool draws two lines connecting these points (*q*1 and *q*3). The user places points S1 and S2 at the ends of the scale bar. **F** Collected measurements from this image are as follows: Length is the distance from L1 to L2. Asymmetry is the ratio of the larger distance among *q*1 and *q*2 to the smaller. Angle of curvature is calculated as the angle formed by points L1, L2 and the midpoint of *q*2. Width is the longest distance between *q*1, *q*2, and *q*3. Aspect ratio is the ratio of length to width. See Table 2 for additional details.

### 3.5 Assessing the accuracy of image measuring software

To examine the possible interactions between shape parameters and the accuracy of the image measuring software, an array of 24 egg silhouettes were simulated with combinations of known parameter values (Figure 3). Each of these eggs was measured five times with the custom image measurement tool to calculate aspect ratio, asymmetry, and the angle of curvature (Table 3).

**Figure 3:**
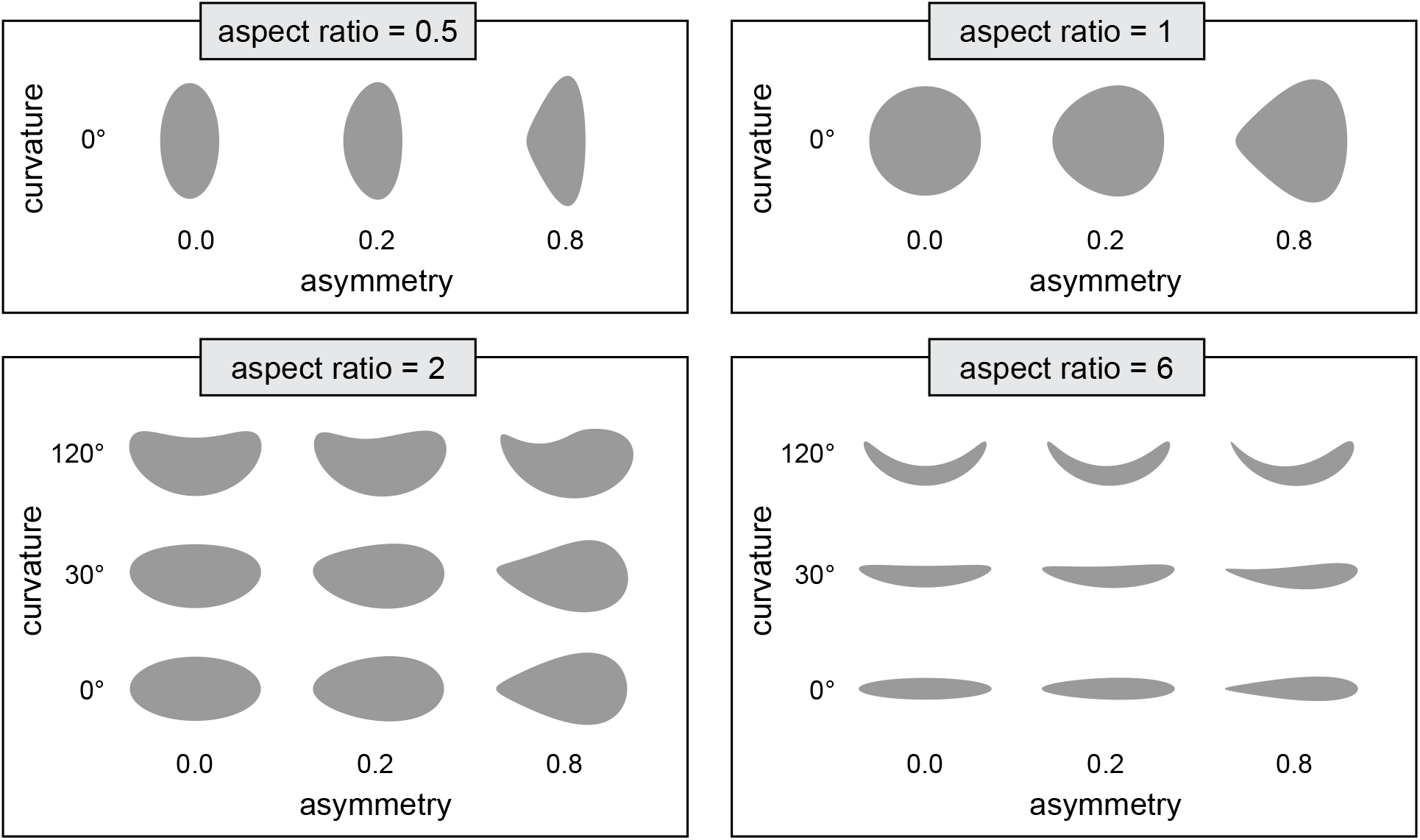
Assessing the accuracy of the egg image measuring software. Simulated egg silhouettes with known combinations of shape parameter values used to assess accuracy of image measurement software. Each egg was remeasured five times using the image measurement software and the results are reported in Table 3.

### 3.6 Calculating final and transformed values

Following data extraction from text and image sources, final values (e.g. volume, aspect ratio) were calculated. Egg length, width, breadth, volume, aspect ratio were log10 transformed, and egg curvature and asymmetry were square root transformed. For entries that had both a text description of egg size as well as an image with a scale bar, the text description was used in the final calculations. Both the raw and processed final database are freely available for download (Dryad https://datadryad.org/review?doi=doi:10.5061/dryad.pv40d2r).

### 3.7 Cross-referencing entries with taxonomic and genetic databases

Taxonomic names parsed from the literature occasionally contained errors, including published typographical errors and optical character recognition errors. These errors needed to be corrected and the taxonomic names also had to be reconciled with currently accepted taxonomy in order to link egg morphology data with other data sources (e.g. published phylogenies). To address these issues, we developed a tool called TaxReformer (https://github.com/brunoasm/TaxReformer, commit 1831a11) that searches the Global Names Architecture (GN)^25,26^, Open Tree Taxonomy (OTT)^27,28^, and Global Biodiversity Information Facility (GBIF)^29^ databases, taking advantage of the strengths of each database. For the taxa included in the insect egg database, GN had the most effective fuzzy matching algorithm and broadest database. OTT provided a better control of the context of each taxonomic query, enabling one to search names only among insects and avoiding homonyms in kingdoms regulated by different codes of nomenclature. OTT’s fuzzy matching algorithm, however, often returned matches to the correct species name but wrong genus name with a high confidence score. OTT and GBIF both contain information about higher taxonomy, which is not standardized in records obtained from GN.

Names obtained from the literature were first parsed with Global Names Parser v. 0.3.1^30^ to obtain genus and species name in canonical forms. The full species name was then used to search in GN with fuzzy matching to allow for correction of optical character recognition errors. If a match to a species or genus was found, the matched name was recorded and then searched in OTT to obtain higher taxonomy and identifier numbers from OTT and the National Center for Biotechnology Information. If the name was not found in OTT, higher taxonomy was alternatively obtained from GBIF. In all cases, if databases contained information about synonyms, the currently accepted name for each taxon was retrieved.

### 3.8 Assessing intraspecific variation

We assessed intraspecific variation in egg size descriptions using four methods:
First, for database entries that reported egg size variation (e.g. egg descriptions that included a range of egg length or an average egg length with deviation), the percent difference in egg size was calculated as follows: for egg descriptions recorded as ranges, percent difference was calculated as 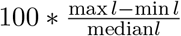; for egg descriptions recorded as average and deviations, percent difference was calculated as 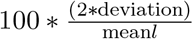.

Second, independent observations of a single species were identified as two entries for the same species that differed in the calculated volume by more than 1.0 * 10^−5^ mm^3^. This excluded entries that were repeated publications of the same description, such as an observation repeated in a subsequent review (Table 1). The percent difference in egg length was calculated as 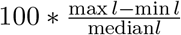.

Third, for entries that had both a text description of egg length as well as a published image with a scale bar, the difference in the reported egg length and our re-measurement of the image was assessed. The percent difference between these two measurements was calculated as 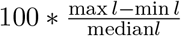.

Fourth, for eggs that were measured as triaxial ellipsoids (length, width, and breadth measured all separately), the percent difference was calculated from the change in egg volume if the egg had been assumed to be a rotationally symmetric ellipsoid 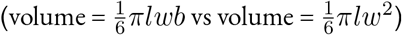. Given that more eggs are likely triaxial ellipsoids than are reported in the egg database, this metric gives insight into the variation in egg volume that might be masked when only two dimensions are reported.

### 3.9 Assessing the precision of entries

The distribution of precision in the insect egg database was assessed using two metrics. First, the number of decimal places used in the length measurement was calculated for each database entry from a base of millimeters (e.g. ‘1 mm’ has 0 decimal places, while ‘1.00 mm’ has 2 decimal places). Second, the relative precision of each measurement was calculated by dividing the total length of the egg by the smallest unit used to measure it, and multiplying this value by 100. This gives the percent of egg length captured by the unit of measurement (i.e. an egg measured as 1.00 mm was measured within 1% of egg length).

### 3.10 Assessing the phylogenetic sampling

The phylogenetic coverage of the insect egg database was assessed by comparing the number of egg entries for a taxonomic rank to the number of species in that rank, estimated by the number of tips in the Open Tree of Life^28^. This assay was performed for all extant hexapod orders and for all insect families in the insect egg database.

## 4 Code availability

All code used to generate the insect egg database as well as reproduce the tables and plots shown here is made freely available. Python code used to compile the database and extract text information from text sources, as well as the R code used to convert the raw database to the final database and to generate the tables and figures shown here is available at https://github.com/shchurch/Insect_Egg_Evolution. Python code used to measure published images of eggs is available at https://github.com/sdonoughe/Insect_Egg_Image_Parser, and python code to cross-reference the egg database with taxonomic tools is available at https://github.com/brunoasm/TaxReformer. Statistical analyses were performed using R version 3.4.2^31^.

## 5 Data records

The final data files include the raw database in tab delimited format, which includes all values extracted from the text and images, as well as the final database in tab delimited format. The code to convert the raw database to the final database is located in https://github.com/shchurch/Insect_Egg_Evolution, directory ‘analyze_data’. Additionally, all data files have been uploaded to Dryad https://datadryad.org/review?doi=doi:10.5061/dryad.pv40d2r.

## 6 Technical validation

The accuracy of the image measuring software was assessed using an array of 24 simulated egg silhouettes with known combinations of parameter values (Figure 4). We found that as the actual angle of curvature increases, the difference between the actual and measured values increases (that is, the software underestimates the angle of curvature), and this difference is larger in eggs with lower aspect ratio and higher asymmetry (Table 3). As the actual asymmetry increases the variance in measured asymmetry increases, and in eggs with low aspect ratio this results in an overestimation of asymmetry. As the actual aspect ratio increases, the software overestimates the total aspect ratio by up to 0.75 (12.5% of the total aspect ratio). Given these results we removed eggs in the top 0.1 percentile of values for asymmetry and aspect ratio when creating the final database.

Intraspecific variation in insect egg size was assessed using four metrics (see Methods section “Assessing intraspecific variation”). The first two describe the percent difference in egg size reported in the literature, either as variation recorded in an egg description (Figure 4A), or as variation recorded across multiple independent observations of eggs from the same species (Figure 4B). In both cases the percent difference in egg length averaged 10% and ranged from 1% to 100% (i.e., for an insect species with an average egg length of 1 mm, it was common to observe eggs from 0.9 to 1.1 mm and occasional outliers at 0.5 and 2 mm.

Additionally we re-measured published images of eggs and calculated the percent difference between our measurements and the text description (Figure 4C). The variation between observations of the same species was consistent with the reported intraspecific variation (average around 10%).

Although the majority of eggs in the database are described as rotationally symmetric ellipsoids (Table 1), for a few clades of insects it is common to measure eggs as triaxial ellipsoids, with length, width, and breadth measured separately (Table 2). Calculating the egg volume using two different methods — one taking into account breadth, and the other assuming rotational symmetry — showed that the percent difference in calculated volume ranges between 10% and 100% (Figure 4D). Eggs from additional clades might be more accurately modeled as triaxial ellipsoids than currently reported in the literature, but this percent difference likely represents the upper range of the error in volume, because the clades typically measured as triaxial ellipsoids are those that are most obviously flattened along one axis.

The text descriptions in the insect egg database were extracted from a diverse set of sources published over hundreds of years, and the precision used to measure eggs varies across these sources (Figure 4). Most entomologists measured eggs in tenths or hundredths of a millimeter (Figure 4E). In terms of the total length of the egg, most measurements in the database are precise to within 1% to 10% (Figure 4F). Given that intraspecific variation is also around 10% of total egg length, it is likely that some of this variation is due to measurement error.

The egg database contains descriptions of eggs from every insect order and from hundreds of insect families (Table 1). Given that the number of species varies greatly across taxonomic ranks we assessed the phylogenetic coverage of the egg database (Figure 4G, H). We found that families and orders with the highest number of estimated species are represented by the greatest number of entries in the egg database. Additionally, most families in the egg database have more than 1 entry per 100 species.

There are several orders represented in the database by fewer than ten entries (Figure 4H). We suggest that this is likely due in part to idiosyncracies of the entomological research for certain clades. For example, although many descriptions of mantis and cockroach oothecae exist, measurements or images of individual eggs within the oothecae are rare in the published literature, which leaves these groups undersampled for propagule size in the literature. The orders with the lowest representation—Trichoptera, Psocoptera, and Zygentoma—are potentially rich new datasets to target for future study.

**Figure 4:**
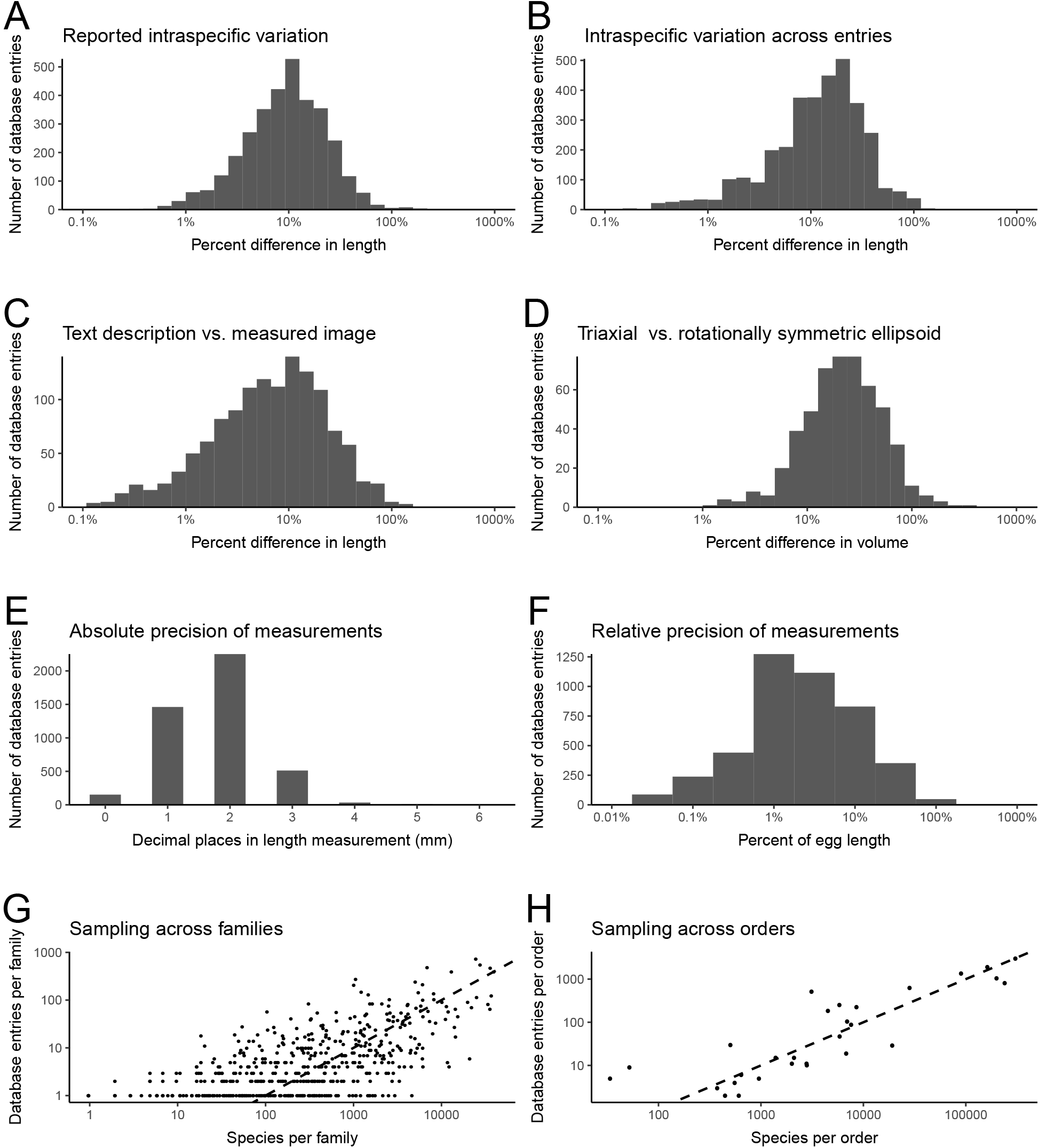
Assessing intraspecific variation, precision, and sampling within the insect egg database. **A**, The distribution of the percent difference between the largest and smallest egg length reported for a species within a publication. **B**, The distribution of the percent difference between the largest and smallest egg length reported for a species across different publications. **C**, The distribution of the percent difference between the largest and smallest egg length, comparing the reported length and the remeasured image from the same publication. **D**, The distribution of the percent difference between the largest and smallest egg volume, measured as triaxial ellipsoids (length, width, and breadth) vs. rotationally symmetric ellipsoids (length and width). **E**, The distribution of the absolute precision of each measurement (decimal places in the egg length measurement in millimeters). **F**, The distribution of the relative precision of each measurement (percent of egg length of the smallest unit used to measure insect egg length). **H**, A comparison of the number of database entries to the number of species estimated in every family present in the insect egg database. **I**, A comparison of the number of database entries to the number of species estimated in every extant insect order. In **H-I** the dotted line shows an arbitrary standard of 1 entry per 100 estimated species.

## Supporting information

## 7 Acknowledgements

This work was supported by the National Science Foundation (NSF) Grant No. IOS-1257217 to CGE, NSF Graduate Research Fellowship No. DGE1745303 to SHC, and by a Jorge Paulo Lemann Fellowship to BdM from Harvard University. We acknowledge Jordan Hoffman and Casey W. Dunn for initial code advice and troubleshooting. We thank the Extavour lab and Brian Farrell for discussion, and Arpita Kulkarni, Angela de Pace, Benjamin Goulet, and Tarun Kumar for suggestions on initial versions of this manuscript. We acknowledge the Ernst Mayr Library at the Museum of Comparative Zoology at Harvard, and specifically Mary Sears, for countless hours of support in gathering the references used in this study.

## 8 Author contributions

SHC and SD wrote all code to parse egg descriptions from the literature, and contributed equally to database creation, study design, writing, and figure preparation. SHC wrote code to manipulate the database and perform statistical analyses. SD wrote code to measure published images. BdM wrote code to correct taxonomic information. BdM and CGE contributed to study design, interpretation, and writing.

